# Single-cell characterization of neovascularization using hiPSC-derived endothelial cells in a 3D microenvironment

**DOI:** 10.1101/2022.02.15.480506

**Authors:** Simon Rosowski, Caroline Brähler, Maren Marder, Misao Akishiba, Alina Platen, Siegfried Ussar, Fabian Theis, Sandra Wiedenmann, Matthias Meier

## Abstract

The formation of vascular structures is fundamental for *in vitro* tissue engineering. Vascularization can enable the nutrient supply within larger structures and increase transplantation efficiency, which are currently limiting factors in organoid research. We differentiated human induced pluripotent stem cells toward endothelial cells in 3D suspension culture. To investigate *in vitro* neovascularization and various 3D microenvironmental approaches, we designed a comprehensive single-cell transcriptomic study. Time-resolved single-cell transcriptomics of the endothelial and co-evolving mural cells gave insights into cell type development, stability, and plasticity. Transfer to a 3D hydrogel microenvironment induced neovascularization and facilitated tracing of sprouting, coalescing, and tubulogenic endothelial cells states. During maturation, we monitored two pericyte subtypes evolving of mural cells. Profiling cell-cell interactions between pericytes and endothelial cells confirmed *in vivo* angiogenic signaling and emphasized new cytokine signals during tubulogenesis. Our data, analyses, and results provide an *in vitro* roadmap to guide vascularization in future tissue engineering.

## Introduction

Endothelial cells (ECs) form the inner luminal epithelium of vascular structures, including arteries, veins, and lymphatic vessels^1^. Apart from core nutrient and oxygen transport functions, the vascular system enables immune cell trafficking, vasomotor tone, and wound healing^2^. To fulfill the different homeostatic functions, ECs exhibit a high degree of plasticity to form differently sized fenestrae and branched vessel structures, recruit organ-specific scaffold cells, or remodel the extracellular matrix composition of the vascular bed^3,4^. The developmental factors determining EC formation, maturation, and specification are not fully revealed yet.

*In vitro* cell culture systems with patient and stem cell-derived ECs have become indispensable for vascular research to overcome the paucity of longitudinal studies in patients, reducing the biological complexity. In particular, stem cell-derived ECs are of interest because they provide access to the early development stage of vascular structures involving cell type formation and specification^5^. *In vivo*, ECs evolve from mesoderm-derived progenitors (angioblasts) in response to FGF2 and VEGFA signals from the adjacent visceral endoderm^6^. Stem cell differentiation protocols recapitulate *in vivo* development by first inducing mesodermal progenitor cells with BMP4 and GSK3β inhibitors^7^, and subsequently ECs by adding VEGFA^8^. Several variations of the differentiation protocols have been reported^9^, where the differences between the resulting ECs remain unresolved. Stem cell-derived ECs do not show a direct correlation with any organ-specific ECs and it remains unclear which developmental stage they reflect^10^. To closely resemble the *in vivo* microenvironment in a dish, 3D stem cell culture formats were introduced, which enable stem cell-derived ECs to form tubes^11^. Moreover, stem cell-derived ECs can fully self-assemble into blood vessels in an organoid-like format, including mural cells, i.e., smooth muscle and pericytes lining the outer surface of the vessel endothelium^12^. In all culture formats, mural cells co-evolve during EC differentiation^13^, which implies inherent strong cell-cell communication during differentiation and maturation.

During embryonic development, immature ECs coalesce and undergo tube formation to produce a vascular plexus^14^, which further differentiates into the specific vessel types. Single-cell transcriptomic analysis of the *in vivo* neovascularization process was performed by extracting cells from laser-induced choroid lesions in mice^15^. The data depicted a highly heterogeneous process with multiple transcriptomic EC stages and types, including phalanx and tip cells, with associated distinctive metabolic profiles. Comparable single-cell transcriptomic analyses in a less complex *in vitro* microenvironment with stem cell-derived ECs is not available but would add fundamental knowledge on the formation of vascular structures.

In this study, we used single-cell transcriptomics to sequentially investigate the development and neovascularization of human induced pluripotent stem cells (hiPSCs) in an *in vitro* 3D microenvironment. In the first step, we used single-cell transcriptomics to explore the differentiation trajectory of co-evolving ECs and mural cells in a 3D suspension cell culture format. Comparison of single-cell transcriptomics of ECs, evolved of a monolayer and 3D suspension culture, revealed differences in ECM gene expression and optimal differentiation parameters. In the second step, the single-cell transcriptomics approach was used to analyze the neovascularization in the heterogeneous 3D suspension culture upon transfer into a hydrogel culture. Ligand-target links between ECs and subcellular pericyte populations were predicted from the single cell transcriptomes during vessel maturation. Our data and analysis provided the resources for future EC specification studies and tissue engineering approaches.

## Results

### Single-cell analysis of endothelial differentiation in a 3D suspension culture

To investigate the differentiation of hiPSCs into ECs in a 3D cell culture format at the single-cell level, we adopted the chemical two-step induction protocol^11,12,16^. Therefore, hiPSCs were differentiated towards the mesoderm germ layer and EC development was induced in the second step (**Fig 1a**). In suspension, the 3D cell cultures were stable over the differentiation and grew from a diameter of 150 µm to 300 µm. While hiPSC-derived aggregates exhibited a uniform spheroidal shape, aggregates from day four showed a more prolate shape. On day nine, 33.2 % of the cells expressed the endothelial marker *CD31* (*PECAM1*), with a standard variation of 5.3% over three biological repeats (**Fig. S1a**). To reconstruct EC development in the 3D suspension culture and define time-resolved cell composition, we performed single-cell mRNA sequencing (scRNA-seq) analysis on 22,192 cells (see **Table S1** for details). Upon Leiden clustering^17^, six cell clusters were identified. With the progression of the differentiation process, the recorded single-cell transcriptomes changed, as indicated by the time-dependent emergence of distinct cell clusters (**Fig. 1b**). All cell clusters could be assigned to cell types by matching known mesodermal and endothelial developmental markers to the differentially expressed genes (DEGs) in the respective cluster (**Fig. 1c**). The cell populations were assigned to pluripotent stem cells (cluster 1), mesenchymo-angioblasts (cluster 2), mural cells (cluster 3), cells undergoing mesenchymal-endothelial transition (MEndoT) (cluster 4), angioblasts (cluster 5), and epithelial cells (cluster 6). At the start of differentiation (day 0), the cell population consisted of homogenous undifferentiated hiPSCs, where over 96% of the cells expressed the pluripotency markers *OCT4, SOX2*, and *NANOG*. Cells assigned as mesenchymo-angioblasts appeared on day three of differentiation and expressed markers for the lateral plate mesoderm^18^, including *HAND1, MESP1*, and *APLNR*. Mural cells observed on day six of differentiation showed reduced *HAND1* expression level, while smooth muscle marker *ACTA2*, pericyte marker *PDGFRB*, and mesenchymal marker *COL1A1* were consistently expressed. MEndoT cells in cluster 4 expressed the same markers but also included the endothelial markers *PECAM1, CDH5*, and *ESM1*. Only a small fraction (0.9 %) of epithelial cells was observed (**Fig. 1d**). Immunohistochemical staining of 3D suspension cell cultures with the markers *PECAM1* and *PDFGRB* showed de-mixing of the two cell populations; however, there was no induction of vessel formation, supporting the angioblast cell state (**Fig. S1e**). To test the robustness of the differentiation approach at the single-cell level, we sequenced the cells from day nine of two independent differentiation experiments. In both cases, a bimodal distribution of angioblasts and mural cells was observed, with comparable distribution numbers (**Fig. S1b** and **c**).

**Fig. 1.**
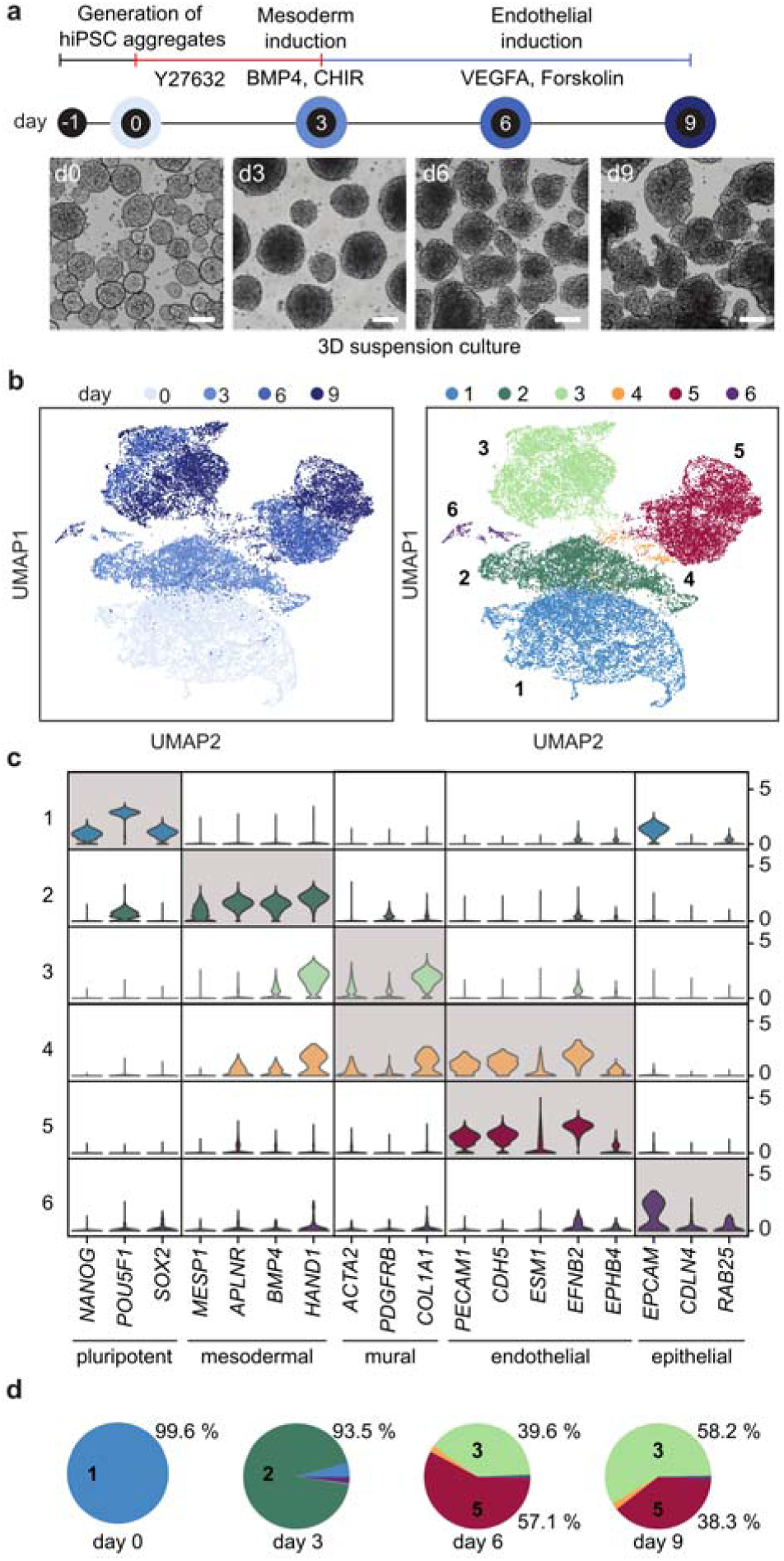
Single-cell transcriptomics reveals the differentiation trajectory of hiPSCs into endothelial cells in 3D suspension culture. **a**, Schematic of the endothelial differentiation timeline with sampling points and chemical induction protocol. Bright-field images show representative 3D suspension cultures at the corresponding time point. Scale: 100 µm. **b**, UMAP plot of the single-cell transcriptomes. Left: Light to dark blue denotes the time points of sampling. Right: Six unique cell clusters were identified during the endothelial differentiation, 1: hiPSCs, 2: mesenchymo-angioblasts, 3: mural cells, 4: MEndoT, 5: angioblasts, 6: epithelial cells. **c**, Violin plot shows the cluster expression levels of differentially expressed genes for the six cell clusters and the commonly used cell markers for cell type assignment. **d**, Cell type distribution analysis along the differentiation trajectory.

### Time trajectory of endothelial development

To resolve the time-dependent relationships of the cell clusters, we performed a dynamic RNA velocity analysis^19^ and identified the dynamic driver genes (DDG) in endothelial differentiation^20^. For this, we first calculated the latent time based on the balance of spliced and unspliced RNA transcripts within the single-cell transcriptomes (**Fig. 2a**). Indeed, the theoretical latent time matched true chronological differentiation times (compared to **Fig. 1a**). The corresponding RNA velocity streamlines indicate two differentiation routes. The differentiation route for angioblasts and mural cells evolved from mesenchymo-angioblasts. Also, velocity streamlines between mural and angioblasts via the intermediate cells undergoing MEndoT indicated a relevant degree of plasticity of the cell types at this stage of development. Evaluation of cell cycle states showed that angioblasts were entirely in the G1 phase, whereas approximately 50% of the mural cells were in G2 and S and thus proliferating (**Fig. S1d)**. Subsequently, we plotted the DDGs along the latent time to trace the central genes for the development of angioblasts in 3D (**Fig. 2b**). Similar to the DEG analysis, cluster-specific DDGs were identified. The list of DDGs for each cell cluster is given in **SI File**. The top DDGs for the mural cell progenitors were involved in cell migration, attraction, or repulsion (*UNC5C, SLIT3*, and *TGFB2*)^21,22^. For ECs, known developmental genes of vasculogenesis were upregulated, including the *VEGF* receptors (*KDR, FLT1*) and the interacting receptors *TIE1* and *TEK*^23,24^. To infer the transcription factors (TFs) controlling the development of angioblasts and mural cell progenitors, a transcription factor enrichment analysis (TFEA) was performed on the DDGs (**Fig. 2c**). The highest-ranked TFs for EC development were *BCL6B*^25^, *ETS1*^26^, *ELK3*^27^, *ERG*^28^, and members of the SOX family. All of them are reportedly associated with the process of early vasculogenesis with context-dependent function but integrate the VEGFA and Notch signaling pathways^25^. Notably, the extracted TFs are only responsible for the development of the two identified cell types, but not for neovascularization since images of sections of 3D cell cultures taken on days six and nine showed no vessel organization (**Fig. S1e**). TFEA of the DDGs for the mural cell progenitors revealed *TBX18, CENPA*, and *HAND2* as the top regulatory TFs^29^. In particular, *TBX18* has been recently identified in mice to be selectively expressed by all pericytes and vascular smooth muscle cells within the retina, brain, heart, skeletal muscle, and adipose fat depots^30^. It has to be considered, that the velocity analysis with spare time points has to be understood as transcriptional correlation rather than real dynamics. Nevertheless, turn-on and -off of genes derived from the velocity analysis match with the expectation, for example for the cell cycle regulator *HMGA2*. In mural cells *HMGA2* is in the turned-on state, whereas in angioblasts in the turned-off state but still detectable.

**Fig. 2.**
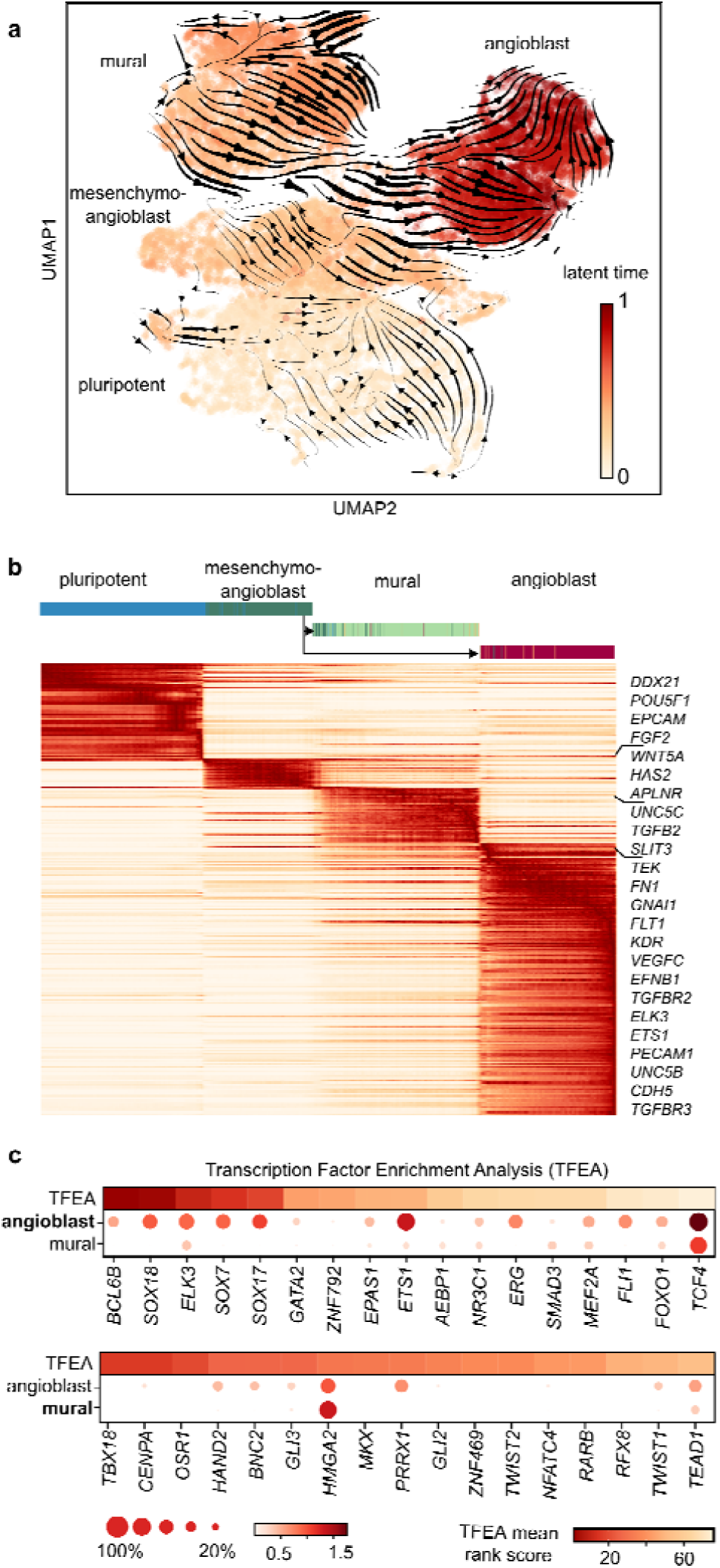
Transcriptomic dynamics predict the differentiation path for endothelial and mural cells. **a**, UMAP cluster plot colored with the latent time calculated based on RNA velocity analysis. The velocity streamlines are given by the black arrows. **b**, The top 300 DDGs sorted according to their likelihood scores and latent time (see **SI File**). **c**, TFEA on the cluster-specific and unique DDGs for angioblasts and mural cells. The dot plots show the gene expression level in the respective single-cell cluster, whereas the upper heat-colored bar shows the TFEA score. Color intensity and dot size denote the normalized cluster mean expression and the fraction of cells expressing the gene, respectively.

### Comparison of endothelial differentiation in 3D versus 2D cell culture formats

In the next step, we sought to compare the *in vitro* development of ECs in 3D suspension culture to a previously performed stem cell differentiation approach with an adherent 2D cell culture on the single cell level^13^. Chemical compounds for the endothelial induction protocol were the same; however, minor concentration differences of the individual compounds existed (**Table S2**). Single-cell transcriptomic trajectories of the 3D and 2D differentiation approaches differed with respect to the marker expression profiles of intermediate and evolving EC and mural cell types (**Fig. 3a**). In the 3D suspension culture, the mesenchymo-angioblast markers (*APLNR, KDR*, and *PDGFRA*) were expressed strongly. In the 2D cell culture, the markers were either lowly expressed or even absent (**Fig. S2a**). To quantify the differences, we combined the two scRNAs datasets. Of the 14,383 genes, 682 and 652 showed expression level differences with a p-value lower than 10^−100^, in the 2D and 3D cell cultures, respectively (**Fig. 3b**). Expectedly, the expression patterns of the ECM genes were distinct in different culture formats. Within the 2D cell culture format, ECs exhibited a strong collagen phenotype with high expression levels of basal lamina proteins, such as *COL4A1/2, COL6A2*, or *COL18A1* (**Fig. 3c**). In 3D cell culture, ECs expressed hyaluronic acid and the corresponding binding proteins. Further, a GO term analysis of the DEG showed that ECs in the 2D cell culture expressed genes associated with migration and motility while Rap signaling was enriched in 3D (**Fig. S2d** and **e**). In contrast, angioblasts within the 3D suspension culture upregulated cell-cell interaction and actin remodeling genes, such as *CLD5, DOCK4*, and *CTNNB1* (Wnt signaling)^31^, and *RAP1B*^32^, *RAPGEF5*^33^, and *RASIP1*^34^ (Rap1 signaling), respectively.

**Fig. 3.**
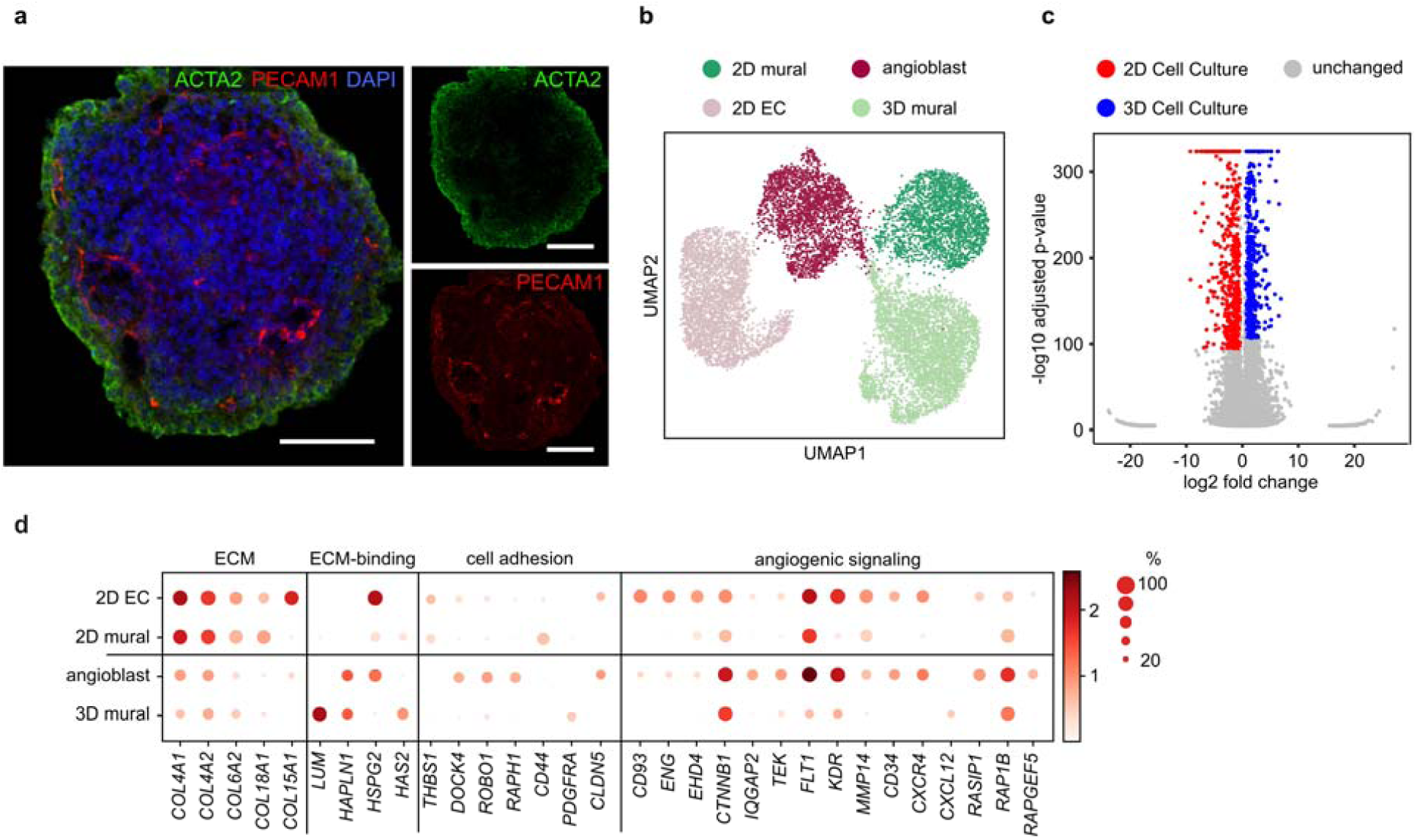
Comparison of EC differentiation in 2D and 3D cell culture formats. **a**, Immunofluorescence image of an aggregate from 3D suspension culture on day 9 of differentiation. Scale: 100 µm. **b**, UMAP plot of single cell transcriptomes of iPSCs derived endothelial cells differentiated on a 2D adhesion monolayer (5267 cells) and as 3D suspension culture (6492 cells). **c**, Volcano plot representing differentially expressed genes between ECs differentiated in 2D and 3D cell cultures. **d**, Representative different expressed genes with assigned cellular functions and biological processes. Color intensity and dot size denote for the normalized cluster mean expression and fraction of cells expression the corresponding gene, respectively.

While mural cells and ECs in the 2D cell culture format were proliferating, within the 3D cell culture, only the mural cells were cycling (**Fig. S2b**). This raised the question of whether the separation of the ECs from the mural cells or the transfer to a 2D culture format reinstates the proliferation of ECs. Indeed, FACS-sorted PECAM1^+^ ECs from the 3D suspension culture proliferated and could be passaged over six generations upon plating on a culture dish. Comparison between sorted PECAM1^+^ cells within a 3D and 2D cell culture was not possible due to the observation that ECs alone did not form a stable aggregate in suspension culture. However, the transfer of the unsorted mixed 3D aggregates to a 2D cell culture format on day six of differentiation led to the proliferation of ECs, demonstrating that growth arrest is associated with the 3D suspension culture format^35^.

A further important question for cell type engineering was whether sorted ECs and mural cells maintained cell type stability after sorting. To evaluate this aspect, we investigated the ratio of PECAM1^+^ and PDGFRB^+^ cells within sorted and unsorted 3D suspension cultures. Within sorted PECAM1^+^/PDGFRB^-^ cells cultures no PDGFRB^+^ cells could be detected after six days of culturing in a monolayer culture format under EC differentiation medium (**Fig. S7b** and **c**). In contrast, PDGFRB^+^ PECAM1^+^ sorted cells lost the PDGFRB expression under the same conditions. Non-sorted cell cultures the fraction of PECAM1^+^ /PDGFRB^-^ cells increased to 72%, where the fraction of PDGFRB^+^ cells also decreased to 10% after six days of culturing in a monolayer. This finding contrasted with that in the 3D suspension culture, where the PDGFRB^+^ cell fraction was unchanged within the same culturing interval. In conclusion, cell-cell signals between angioblasts and mural cells lead to stabilization of the mural cell fraction.

### Single-cell transcriptomics of in vitro sprouting and coalescing ECs

Stem cell-derived angioblasts generated under comparable conditions can be induced by sprouting^12^. We induced endothelial sprouting by transferring the 3D suspension culture into Matrigel (**Fig. 4a**). Sprout formation was observed within the first 12 h (**Fig. S4d**). Single-cell transcriptomes of the sprouting Matrigel culture were determined 48 h after transfer and compared to single-cell transcriptomes of cells within 3D suspension cultures kept to the same day of differentiation. ECs formed two transcriptomic subclusters within the Matrigel culture, which were separated from ECs in the 3D suspension culture (**Fig. 4b**). In addition, mural cells formed two transcriptomic subclusters, where one cluster overlapped with the transcriptomic state of the mural cells in the 3D suspension culture. Transcriptomes of MEndoT cells clustered with their analogs in 3D suspension culture (**Fig. 4c**). The expression levels of general cell type markers are shown with cluster assignments in **Fig. S3a**. A corresponding velocity analysis with a calculated latent time of the combined datasets indicated that the transcriptional states of ECs observed in the Matrigel culture evolved from the suspension culture, similar to the mural cells (**Fig. 4d**). Further, the streamlines of the velocity analysis indicated that MEndoT cells developed from the mural cells towards EC within the Matrigel microenvironment, which is consistent with the velocity analysis of the 3D suspension culture.

**Fig. 4.**
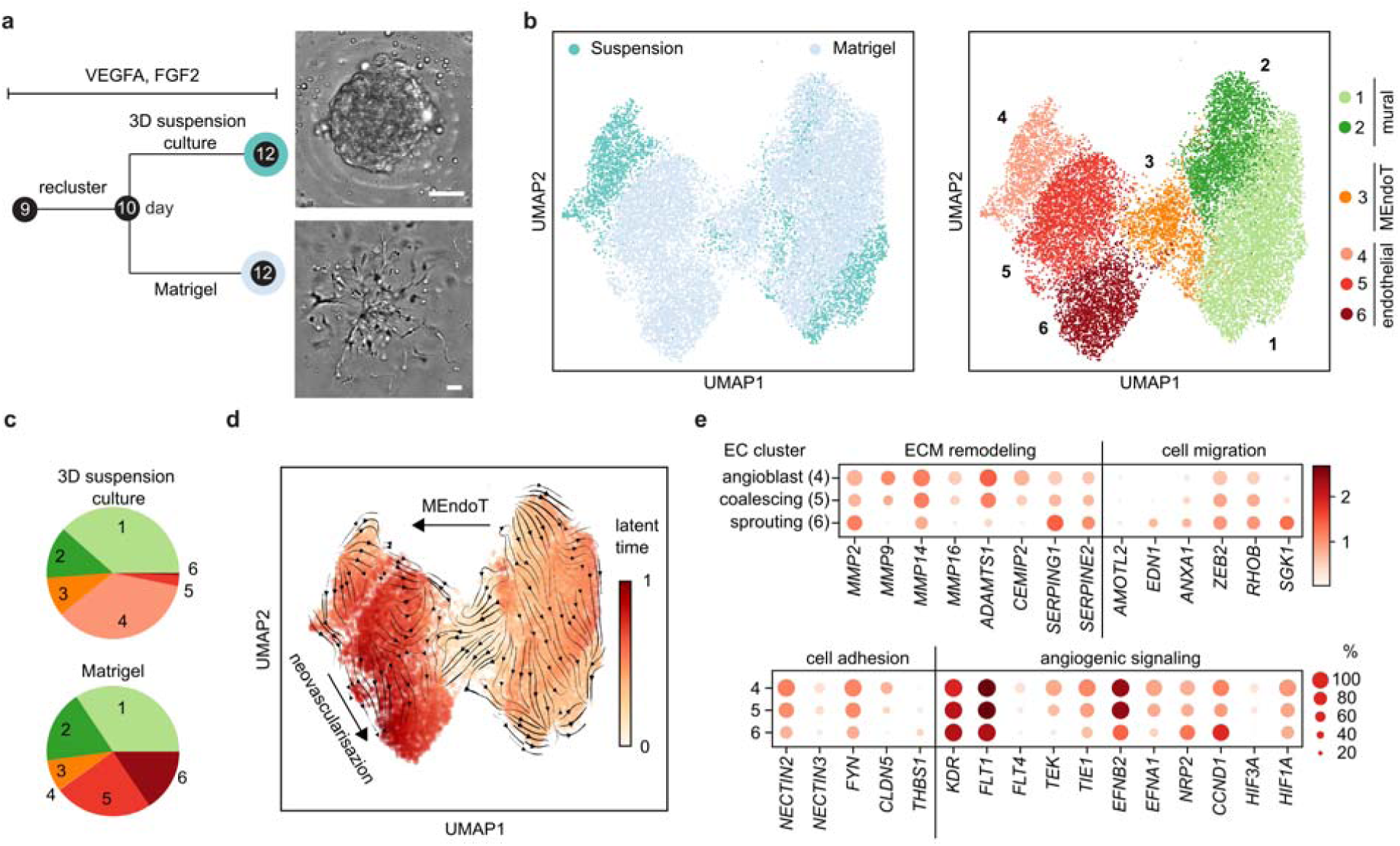
Single-cell analysis of hiPSC-derived ECs undergoing neovascularization in Matrigel. **a**; Experimental overview of the culturing conditions for microvessel formation induction and representative bright-field images of cell culture morphologies. Scale: 50 µm. **b**, UMAP plots containing single-cell transcriptomes of cells from 3D suspension and Matrigel cultures. The color code denotes conditions (left) and Leiden cell clusters (right; 1 and 2: mural cells, 3: MEndoT, 4: angioblasts, 5: EC coalescing, 6: EC sprouting). **c**, Cell type composition of the two culturing conditions is represented as a pie chart. **d**, Velocity analysis of the single-cell transcriptomic data from **a. e**, Expression levels of representative DEGs sorted by function for the three EC transcriptional states. Color intensity and dot size denote the normalized cluster mean expression level and the fraction of cell expression for the corresponding gene, respectively.

For the formation of microvessels, angioblasts within the Matrigel-embedded 3D suspension culture must migrate into the hydrogel and coalesce. To infer which genes drove the processes, we plotted the DEGs related to ECM remodeling, cell migration, cell interaction, and VEGF signaling (**Fig. 4e**). Based on the DEG cluster profiles and corresponding GO term analysis, EC clusters 5 and 6 can be assigned to coalescing (cEC) and sprouting (sEC) endothelial cell states, respectively (**Fig. S2c** and **d**). The transfer to Matrigel increased the expression levels of genes involved in migration within the ECs (e.g., *AMOTL2*^36^, *ANXA1*^37^, or *ZEB2*^38^) and reduced the expression levels of genes involved in cell interactions (*NECTIN2/3* or *CLDN5*). ECM proteins required for the build-up of a basal lamina increased in cECs (*COL4*^39^, *COL18*^40^). The expression levels of integrin changed only slightly between the three EC transcriptional states. Moreover, only the collagen type 1 metalloproteinase (*MMP*) *2* was gradually upregulated upon cell state transversion from angioblasts into cECs, whereas *MMP9* and *MMP14*, which are known to promote motility and tubulogenesis, respectively, were downregulated^41,42^. The most obvious was the downregulation of genes related to the VEGF signaling network within cECs and sECs, except for *KDR* and *NRP2*, which both act on endothelial motility, sprouting, and survival^43,44^. A corresponding TFEA of the DEGs between cEC and sECs showed enrichment of *ELK3, KLF6*, and *SNAI2* (**Fig. S3d**). The latter is a TF well described in the context of the epithelial-mesenchymal transition^45^, which supports the migrating character of the sEC state. Additionally, *ERK* and *KLF6* control cell proliferation, which was indeed reinstated in ECs after Matrigel transfer^46^. This is further supported by the downregulation of Notch signaling genes in the sprouting cell state, including *NOTCH1/4, DLL4*, and *JAG1* (**Fig. S3c**). Notch inhibition in cellular model systems has been shown to induce sprouting, branching, and filopodia induction^47–49^. Furthermore, the mTOR pathway proteins, particularly those of the mTOR complex 2 (RICTOR), were strongly downregulated in sECs (**Fig. S3c**). One downstream target of mTORC2 is the serum and glucocorticoid kinase 1 (*SGK1*), which is the top upregulated gene in sECs, indicating strong metabolic regulation in this motile cell state^50,51^.

### EC maturation in Matrigel

Upon prolonging the culturing time, the vessels grew and branched in the Matrigel microenvironment. To determine the genes activated during vessel maturation, we acquired single-cell transcriptomes of day 18 Matrigel cultures. In addition, we investigated the effect of ascorbic acid (AA) on EC maturation. AA increases the synthesis of the basal laminal protein collagen IV and reduces vessel permeability^52,53^. Bright-field imaging showed that vessel length and branching network were comparable in the presence and absence of AA (**Fig. 5a**). To extract DEGs responsible for EC maturation, single-cell transcriptomes of day 12 and 18 Matrigel cultures (with and without AA) were clustered together (**Fig. 5b**). Day 18 ECs in the presence and absence of AA clustered together, where a low fraction of sprouting and coalescing cells was still observed on day 18 (**Fig. 5c**). The addition of AA reduced the fraction of sEC in the culture by an order of magnitude. The transcriptomes of the mural and MEndoT cells from day 18 converged partially with the transcriptomes of day 12 and formed an additional cluster. The cell-type composition within the Matrigel culture changed from days 12 to 18. While the ratio of ECs to mural cells was balanced on day 12, on day 18, the amount of mural and MEndoT cells increased up to 66 %. In the presence of AA, the fraction of mural cells and MEndoT cells further increased to 85 % (**Fig. 5c**) due to the increased number of proliferating mural cells (**Fig. S5c**). The velocity analyses again indicated that the MEndoT population developed from the mural cells towards ECs (**Fig. 5d**), where day 18 MEndoT cells were further committed to ECs, which is shown by the loss of *PDGFRB* and further mesenchymal markers (**Fig. S4a**). Evaluation of the DEGs between ECs revealed that on day 18, ECs increased the expression levels of cell-cell contact genes as *ICAM*s and *CLDN5* (**Fig. 5e**). Furthermore, the expression of key tubulogenesis genes, such as *RASIP1, RHOB, ELMO1*^54^, and *ARHGAP29*^55^, were upregulated. In addition, *DOCK9* gene expression level increased, which is an *RAC1* activator responsible for vascular lateral branching^56^. Based on the expression pattern, we assigned day 18 ECs to tubulogenic EC (tEC). In line with the velocity analysis, a Pearson correlation of the EC transcriptional states from days 12 and 18 showed that the tubulogenic state was closer to the coalescing than to the sprouting state (**Fig. S4c**). In addition to the changes in the levels of genes controlling the structural change, Notch signaling was upregulated again compared to the sprouting and coalescent states in the forms of *NOTCH1, 4* and *DLL4*. tECs within the +AA sample exhibited a higher latent time within the velocity analysis, indicating a more mature state than without AA. Upon increasing the Leiden cluster parameters, it is possible to separate ECs cultured in the presence and absence of AA; however, DEG change was minimal. Transcription factor analysis of the DEGs from tECs did not reveal differences in coalescing ECs.

**Fig. 5.**
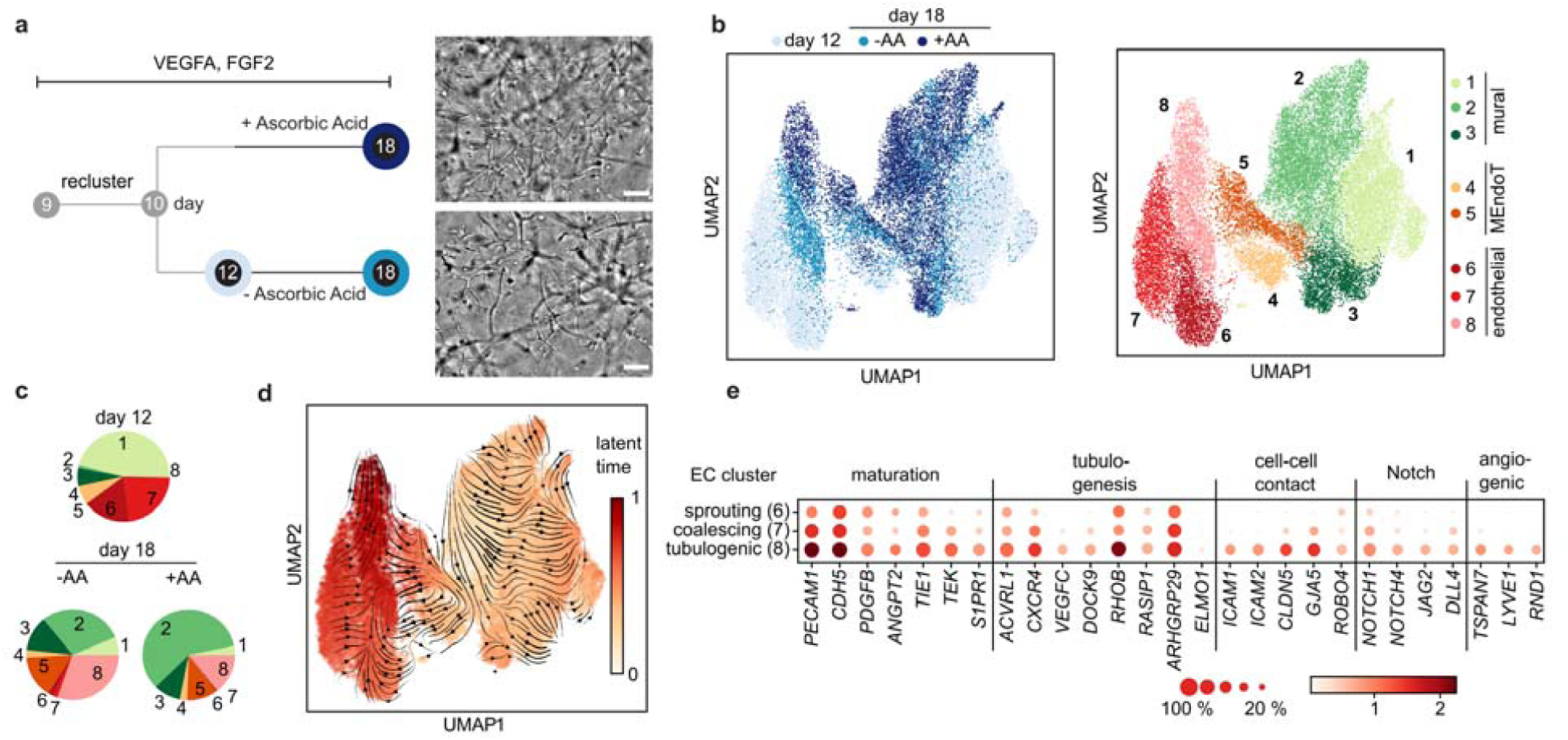
Maturation of endothelial cells in Matrigel in presence and absence of ascorbic acid. **a**, Experimental timeline and applied conditions with corresponding bright-field images of stem cell-derived EC and mural cell cultures in Matrigel on day 18. Scale: 50 µm. **b**, Left: UMAP projection of scRNA-seq data from day 12 and two samples from day 18. Right: UMAP projection colored for the annotated cluster 1: mural cell, 2 and 3: pericytes (P1 and P2), 4 and 5: MEndoT, 6: sprouting ECs, 7: coalescing ECs, 8: tubulogenic ECs. The dataset contains 13159, 6373, and 8684 cells for days 12 and 18 without and with ascorbic acid in the media, respectively. **c**, Cell type composition in the samples. **d**, Velocity analysis of the single-cell transcriptomes, where the latent time is colored on the UMAP plot. **e**, Expression levels of representative DEGs sorted by function for the three EC transcriptional states. Color intensity and dot size denote the normalized cluster mean expression level and fraction of cell expression for the corresponding gene, respectively.

### Mural and Endothelial cell-cell interaction

The single-cell transcriptomes of the microvascular culture on day 18 revealed that mural cells of clusters 2 and 3 (**Fig. 5b**) expressed pericyte markers *NG2, RGS5*, or *NT5E*. Among the DEGs were further genes previously found to be enriched in pericytes, such as *POSTN* and *PDLIM3*. Therefore, we assigned mural clusters 2 and 3 as pericytes, P1 and P2, respectively. While cells of the P1 cluster expressed the transcription factor *FOXF1*, cells of the P2 expressed *GATA4* (**Fig. 6a**), where both found in primary pericytes^57^. Localization of the subcellular pericytes within the Matrigel culture failed either due to the low amount of specific antibodies for the DEGs and cross expression of the marker. Mural cells of cluster 1, which consisted mainly of cells from day 12 and clustered with mural cells of the 3D suspension culture. The higher expression level for the mesenchymal TF *HAND1/2* and *ACTA2*, arguing that mural cells of cluster 1 were in a premature stage. The velocity analysis of the mural cell transcriptomes, however did not show any direction or evolving latent time underlining the plasticity of the cells.

**Fig. 6.**
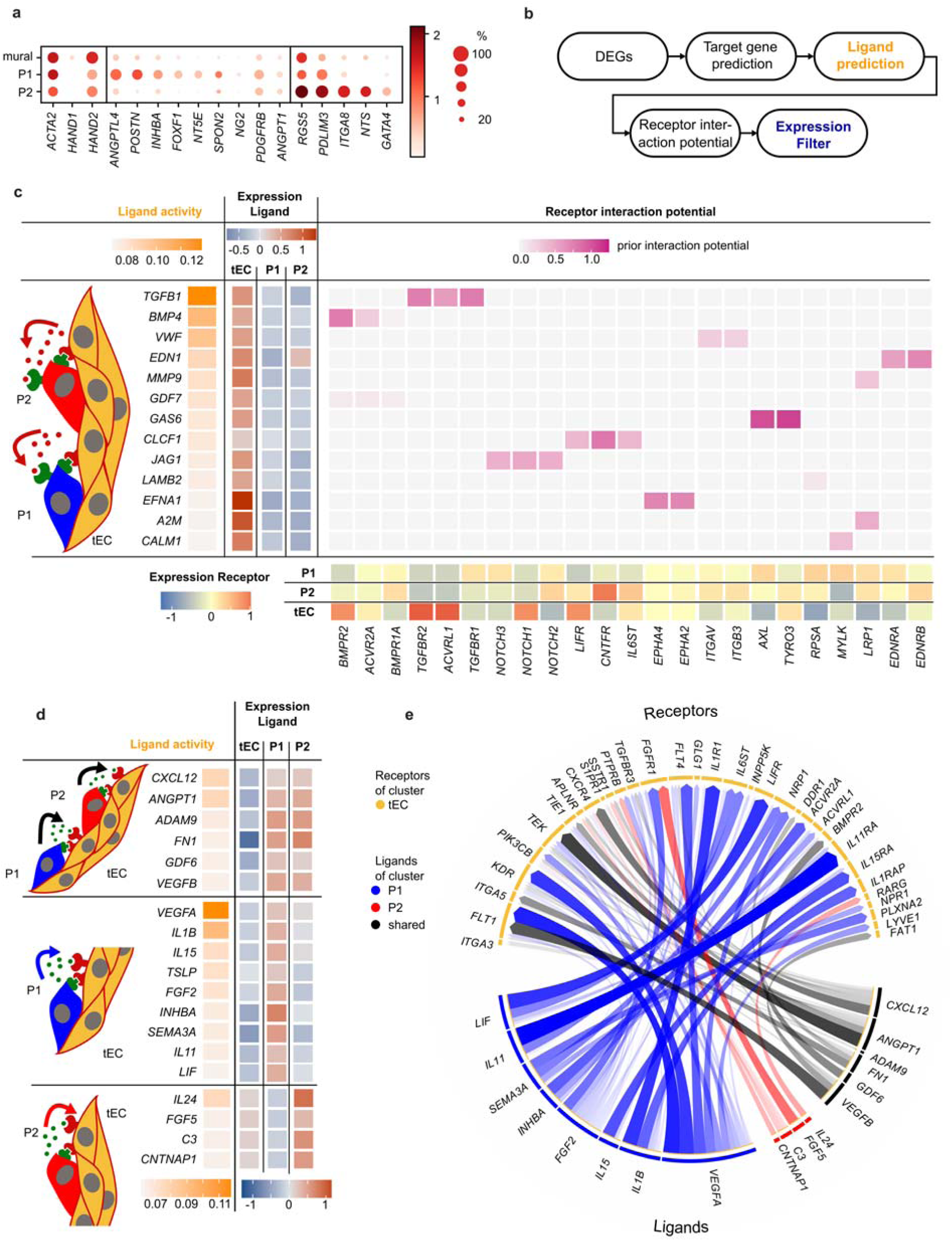
Inferred signaling between endothelial cells and pericytes during vessel maturation from single-cell transcriptomics. **a**, Gene expression analysis of the mural cells cultured in Matrigel up to day 18 of differentiation. The P1 and P2 clusters show the expression of pericyte markers (bold). Color intensity and dot size denote the normalized cluster mean expression level and fraction of cell expression for the corresponding gene, respectively. **b**, NicheNet analysis workflow with the expression filter to infer ligand-receptor interactions. **c**, Ligand-receptor pairs inferred with NicheNet, where single-cell transcriptomes of tEC are used as senders and the combined P1 and P2 pericytes subpopulation as receivers for the analysis. **d** and **e**, Ligand-receptor pairs inferred with NicheNet, where single-cell transcriptomes of the P1 and P2 pericyte subpopulations are used as senders and tEC as receiver cells for the analysis. Overlapping (top rows) and unique pericyte ligands (middle and bottom rows) are listed in table format. Ligand-receptor pairs are presented in the circular chord diagram.

In the last step, we asked whether it is possible to infer cell-cell communication signals leading to vessel maturation from the scRNA datasets of high plastic cell types, stability, reduced proliferation, and migration^58,59^. To trace cell-cell signaling between pericytes and tECs, we performed a ligand-receptor analysis on the single-cell transcriptomic data. Algorithms evaluating only ligand-receptor expression profiles could not recapitulate literature confirmed signals between ECs and pericytes, for example, TGFB1 or VWF (**Fig. S6a and b**) in our dataset. Therefore, we performed a NicheNet analysis^60^, which infers ligand-target links between interacting cells by combining DEG data with prior knowledge of signaling and gene regulatory networks (**Fig. 6b**). NicheNet was applied bidirectionally by investigating ECs as ligand sender and receiver cells. According to the NicheNet results, that is, ligand activities and receptor and target gene interaction potential, were further filtered for gene expression levels to increase the probability of finding relevant interaction links (**Fig. 6c**). For ECs as senders and pericytes as receivers, the algorithm predicted strong ligand activity for *TGFB1, BMP4*, and *VWF*. All three have been reported previously to be central for communication between both cell types^61^. *TGFB1* acts in paracrine and autocrine signaling, while tECs express the *TGFB1* receptor *ALK1* (*ACVRL1*) and pericytes express *ALK5* (*TGFBR1*), which is consistent with current reports using *in vitro* culture systems^62^. The inferred ligand activity for *PDGFB* was low and thus filtered out, although *PDGFB* was highly expressed. This is explained by the fact that we used DEGs of the entire differentiation trajectory to rank ligands, and since *PDGFB* is expressed strongly at all times, the corresponding target genes within the DEG cannot be expected. In turn, the top predicted ligand activities represented signals that occurred during vessel maturation rather than during vessel formation and the pericyte recruitment phase. Individual NicheNet analysis between ECs and P1 or P2 cells did not differ from the combined analysis. In contrast, the NicheNet analysis for P1 or P2 as signal senders and tECs as the receiver showed that P1 and P2 pericytes exhibited shared, as well as individual ligands (**Fig. 6d**). The shared ligands included *CXCL12, ANGPT1*, fibronectin, and *GDF6*, which antagonize VEGF signaling to promote junctional stability and vascular integrity^63,64^. The strongest individual predicted ligand activity of P1 cells was *VEGFA*, followed by a set of cytokines (*IL1b* and *IL15*) and *FGF2*. For P2 cells, the ligands with the strongest individual activity were *IL24* and *FGF5*. The corresponding tEC receptors are shown in **Fig. 6e** and in the detailed table in **Fig. S6c**. All inferred cytokines have proven pro-or anti-angiogenic function in cancer angiogenesis (**SI File**); however, their function during neovascularization and vessel maturation is unknown. The resolved heterogeneity of cytokine ligands between pericytes argues for functional differences between P1 and P2 during vessel maturation.

## Discussion

Here, we investigated the differentiation of hiPSCs into ECs in a 3D suspension culture, and subsequently, the process of neovascularization with the derived ECs and co-evolving mural cells at the single-cell level. We found that the general differentiation trajectory of ECs and co-evolving mural cells in a 3D suspension culture resembled the development of that in a monolayer format. In contrast to the 2D cell culture approach, ECs become quiescent in the 3D suspension culture without entering the neovascularization process. Within the 3D suspension culture the ECM in the surrounding of angioblasts did not contain COL1. Furthermore, COL1 interacting proteins as the *ITGA1* receptor were low expressed in angioblasts. *ITGA1* signaling is essential for *in vivo* angiogenesis and could explain the observed proliferation arrest. COL1 expressed by the mural cells could not compensate the missing COL1 since its deposition was spatially restricted to the mural cells in the 3D suspension culture. This also holds true for anti-angiogenic factors, such as Lumican (LUM), which colocalized only with mural cells (**Fig. S2e**). Cell communication between angioblasts and mural cells exists within the 3D suspension culture, which is exemplified by the finding that upon separation from the ECs, the plastic mural cells lost mesenchymal cell type marker expression of *PDGFRB*. Mesenchymo-angioblast cells express *TGFB1* and *PDGFB*, where the combination of these two factors is used to induce vascular smooth muscle cell development *in vitro* in the same mesodermal precursor cells^16^. Interestingly, stimulation of mesenchymo-angioblasts with ActivinA (INHBA), a member of the TGFB family, and PDGFB led directly to PDGFRB^+^ mural cells as reported before. Additionally to the PDGFRB^+^ cells, we detected in the differentiation approach a fraction of 7% of PECAM1^*+*^ cells on day six and 6% on day nine (**Fig. S7a** and **b**). This demonstrates that EC and mural cells co-evolve in both differentiation directions *in vitro*. Lineage specification on day 6 of differentiation into an arterial tone could be observed when taking the stronger expression level of *EFNB2* into account^65^ (**Fig. 1c**). For scaling EC production with a 3D suspension culture, the transcriptional cell type analysis argues that six days of differentiation is optimal since it shows the largest EC to mural cell ratio.

Transfer of the 3D suspension culture into Matrigel induced sprouting, where the collagen 1 type hydrogel was sufficient but showed a slower growth rate (data omitted). Single-cell transcriptomics untangle coalescence, sprouting, and tubulogenic EC cell states during neovascularization. In comparison to the *in vivo* reported data, after laser ablation in the choroid layer of mouse eyes, the cellular complexity is far lower. In addition to the expected VEGF and Notch signaling pathways, the mTOR pathway is regulated during neovascularization. mTOR kinase controls various processes, when complexed in the mTORC1, it controls predominantly the cell metabolism. The metabolism as for example a switch between OXPHOS and glycolysis for sECs could not be observed (**Fig. S3**), as described for tip cells *in vivo*. However, in sECs, mTORC2 adaptor proteins were downregulated and the downstream effector kinase *SGK1* was strongly upregulated, which controls cell survival during angiogenesis^66^ and EC shape^67^; its ablation led to reduced neovascularization and impaired cell migration^50^. The function and regulation of *SGK1* through mTORC2 are unknown but have become the focus of further investigations.

Vessel structure formation within the hydrogel culture was accompanied by a change in ECM-integrin expression level. While *ITGA2* was upregulated in sEC, a gradual decrease in *ITGA1* accompanied by an increase in *ITGA1, A5*, and *V* was detected along the cEC to tEC cell stages. Blocking collagen-binding *ITGA1* and *A2* with antibodies has been shown to reduce angiogenesis^68^. The EC and mural cell stage-assigned integrin profiles provide new dynamic insights. The addition of AA to the cell culture media increased the expression level and deposition of the central basal lamina protein COL4 (**Fig. S5**). Therefore, the AA-induced maturation of ECs increased the proliferation rate of mural cells. This may be unwanted in long-term cultures during tissue engineering with no proliferative cells. ECs on day 18 of differentiation did not show a specification marker for arterial or any other vascular cell type. The remaining plasticity of the ECs can be once more exemplified by disseminating of the Matrigel culture and reformation of the 3D suspension culture, where corresponding single-cell transcriptomes show the interconversion of the cell states (**Fig. S7e** and **f**). The most compelling results of the single-cell analysis during the vessel maturation phase in a reduced *in vitro* microenvironment were the resolved cell-cell communication signals between the two transcriptional pericyte subpopulations and tECs. Pericytes are recruited by PDGFB signaling to capillary walls to stabilize integrity and tube assembly^58^. The directionality of the

NicheNet ligand-receptor analysis allowed for the resolution of *TGFB1* signals in the sender and receiver cells. Next to the known EC-pericyte signaling factors, the complex cytokine profiles of the pericytes were most obvious. The maturation factor for pericyte formation is deciphered as the CXCR4/CXCL12 signaling axis, but in addition to a variety of chemokines with unreported functions during the neovascularization process^69^. Using the DEGs calculated from transcriptional time trajectories within the sender-receiver model, it is possible to extract cell-cell communication signals at defined processes, as applied in the vessel maturation phase. The present single-cell transcriptomic data and analysis of the vascular structure formation process can be used as a benchmark set for future *in vitro* vascularization approaches, differentiation attempts to alter the specification of ECs, or investigation into disease-specific gene functions.

## Methods

### 2D hiPSC cell culture

HiPSCs were cultured on hESC Matrigel-precoated 6-plates according to manufacturer’s recommendations (Corning) in mTeSR1 medium (Stemcell Technologies) at 5% CO2, 5% O2, and 37°C with daily medium change and split twice a week in a 1:6 ratio using 0.05% Trypsin-EDTA (Sigma).

### 3D suspension culture

For the transfer into a 3D hiPSC cell culture, the medium was aspirated, cells were washed with 2 mL PBS -/- and 500 µL Accutase were added. During incubation at 37 °C for 3 to 7 min, cells detached while adding 2.5 mL mTeSR stopped the reaction. The well was washed with 1 mL mTeSR before centrifugation for 5 min at 200 x g and resuspended in 500 µL mTeSR with Rock inhibitor (1:1000) and 1% Penicillin/Streptomycin (P/S, Thermo). The cells were incubated in ultra-low attachment plates at 37°C and 100 rpm (Fisher).

### 3D suspension culture differentiation to ECs

At day -1 of the differentiation, cells were transferred into low attachment 6 well plates in order to form embryonic bodies (1.5 × 10^6^ cells/well). Differentiation was performed according to^11,16^. Briefly, from day 0 to 3 N2B27 medium supplemented with BMP4 (25 ng/mL) and CHIR (7.5 µM) was used without media exchange. Media was changed daily from day 3 to 7 using StemPro-34 with VEGFA (200 ng/mL) and forskolin (2 µM). Afterward, StemPro-34 with VEGFA (30 ng/mL) and FGF2 (30 ng/mL) was used with exchange after two days.

### 3D microwell chip

For matrix-free cultivation, the microwell chip platform, described by Wiedenmann *et al*., was used to allow the growth of 1196 aggregates in parallel^70^. The chip was coated with 10 % Pluronic F-127 (Sigma) overnight and seeded with approximately 1×10^6^ cells. StemPro-34 with VEGFA (100 ng/mL), FGF2 (100 ng/mL), and FBS (15%) was used for cultivation. The medium was exchanged every second day.

### Hydrogel cell culture

For cultivation in 24-well plates, 100 µL Matrigel were added into each well of the plate that whole ground is covered and incubated at 37°C for 1 h. Aggregates were centrifuged at 800 rpm for 5 min at 4°C, resuspended in 80 µL of Matrigel and spread on top of the first layer. Gel polymerization was done at 37°C inside a container with a bit of ice to allow a slow temperature adjustment. After one hour 0.5 mL media were added. Media composition complies with microwell chip incubation. +AA samples contained 60 µg/mL ascorbic acid. For imaging, the aggregates were embedded in ibidi slides with a number of 10 to 20 per well. The amount of Matrigel and media was adjusted to the smaller volume.

### Fluorescence-activated cell sorting (FACS)

Harvested cells were washed three times with PBS (200 x <u>g</u> for 5 min) and singularized using accutase. Five volumes of FACS buffer (10% FBS in PBS) were added and centrifuged at 300 x g for 5 min. Blocking was done for 20 min with StemPro34 + 10 % FBS on ice. The cell suspension was filtered using a 70 µm nylon cell strainer (Corning). Live dead staining was performed with Trypan Blue while cells were counted. 20 μL of each antibody (FITC Mouse Anti-Human CD31 (BD Pharmingen™, 555445) and PE Mouse Anti-Human CD140b (BD Pharmingen™, 558821)) and 1 µL of violet fluorescent reactive dye (Invitrogen, REF: L34963A, LOT: 2179253) were added per 10^6^ cells and incubated for 30 min at room temperature (RT). After washing once with FACS buffer, cells were resuspended in Tyto buffer (10^6^ cells/mL, MACSQuant Tyto (TM) Running Buffer - Milteniy (Cat. No: 130-107-207; Lot: 5200608355)). The sorting was done on a MACSQuant® Tyto™. For ensuring unspecific binding, FITC Mouse IgG1, κ Isotype Control (BD Pharmingen™, 554679) and PE Mouse IgG2a, κ Isotype Control (BD Pharmingen™, 559319) were used.

### Flow cytometry

Singularized cells were washed once with PBS (centrifugation conditions: 300 x g, 5 min). Cells were fixated at RT for 15 min with 4% PFA in PBS and afterward washed twice with FACS buffer. 10^5^ cells were transferred into a U-bottom-shaped 96-well plate (Greiner) and an antibody (2 µL per 15^5^ cells) diluted in 100 µL FACS buffer was applied per well. After incubation of 30 min at RT, cells were washed twice with FACS buffer. For the measurement, the pellet was resuspended in 200 µL of FACS buffer and transferred through a cell strainer into FACS-tubes (5 mL round-bottom tubes, Coring). Flow cytometry analysis was performed on a MACSQuant® VYB.

### Cell type stability experiments

On day 6 (**Fig. S7c** and **d**), aggregates were harvested and prepared for FACS, ECs (PECAM1^+^, PDGFRB^-^) and VSMCs (PDGFRB^+^, PECAM1^-^) were sorted. 2 × 10^5^ of the sorted cell types and the unsorted mixture were seeded and cultured in 6-well plates coated with fibronectin bovine (LIFE Technologies). Medium composition was equivalent to the 3D differentiation.

### Cryoembedding

For cryoembedding of 3D suspension culture aggregates, 500 µL of 4% PFA were added and incubated for 15 min on ice. The disc was detached from the walls with a needle. After two washing steps with PBS, an incubation with, first, 10%, second, 30% sucrose at RT for 2 h and, third, with a 1:1 mixture of 30% sucrose and OCT medium at 4°C overnight followed. All incubation steps were implemented on a wave shaker at 47 rmp. The medium was replaced by pure OCT and the sample frozen on dry ice. The Slicing was done with a Leica CM1860 cryostat and a thickness of 20 µm.

### Fluorescence imaging

Slides were washed with PBS, permeabilized in PBS with 0.1% Triton for 30 min at RT, washed with 0.2% Tween 20 in PBS (PBST) and blocked with 2% BSA (Proliant) in PBST for 1 h. Antibodies were applied in the concentration as recommended by the manufacturer (specifications in **SI File**) in the blocking solution. After primary and secondary antibody staining, five washing steps with 5 min PBST were performed. Prior to confocal imaging (Zeiss Axio Observer LSM 880), Vectashield® Mounting Medium was added to the sample, it was covered and sealed by a coverslip.

### Image analysis

IF and bright-field images were corrected for brightness and contrast with ImageJ. Z-projection of fluorescence images was performed using maximal intensity. ImageJ version 1.52p was used^71^.

### Sample preparation for scRNA-seq

Matrix-free samples were washed with PBS, resuspended in Accutase and incubated for cell detachment at 37°C for 30 min. The reaction was stopped by adding five volumes of media. Afterward, cells were washed once with PBS. Prior to this process, Matrigel embedded samples were washed with PBS, incubated with 0.5 mL Collagenase/Dispase Solution (1:100 to a final concentration of 1 mg/mL in StemPro34) for around 4 h, until organoids detached from the Matrigel. The enzymatic reaction was stopped with 1 mL Neutralisation Buffer (1%BSA, 1%P/S in DMEM:F12). The single cells were cryo-preserved in DMEM with 10% heat-inactivated FBS (Thermo Fisher Scientific) and 10% DMSO based on a previously described scRNA-seq sample preparation protocol^72^. For sequencing, cryo-preserved cells were thawed in DMEM:F12. RNA libraries were generated using Chromium Single Cell 3’ library and gel bead kit v3.1 (10x Genomics). The amplified cDNA library was sequenced on a NovaSeq 6000 S2 flow cell from Illumina. The sequenced cell numbers can be found in **Table S1**.

### ScRNA-seq data pre-processing

Sequencing raw files were demultiplexed, aligned (reference genome hg38_ensrel97), filtered, barcodes and UMIs counted, and subjected to a quality filter with CellRanger (version 3.0.1, 10xGenomics). The pre-processing and downstream analysis were performed with the package ‘Scanpy API’ in python with default parameters, if not stated differently^73^. First, dead or stressed cells, identified by a percentage of mitochondrial genes higher than 10%, were filtered out. Next, cells with less than 200 and genes expressed in less than three cells were excluded. Afterward, the datasets of different days and experiments were concatenated, normalized to 10^4^ gene counts per cell and log-transformed. Batch effects were corrected using ComBat. Further on, the top 4,000 highly variable genes were used for the downstream analysis. As discussed by Luecken and Theis, we corrected for the total gene counts, percentage of mitochondrial genes, and the cell cycle distribution of S, G2 and M phase to investigate differentiation-dependent changes on the transcriptome level^17^.

### Dimensionality reduction, clustering, and cell-type annotation

The single-cell nearest neighborhood graph was computed with the first 50 principal components and ten nearest neighbors. The cells were clustered with the Leiden algorithm with a resolution of 0.5. For visualization, the dimensionality of the data was reduced using Uniform Manifold Approximation and Projection (UMAP)^74^. For cell-type annotation, 300 DEGs for each of the clusters were calculated by ranking the clusters against all remaining cells with the t-test method (**SI File**). Clusters with proteasome-related genes scored at the top or a significantly reduced gene count were removed from the dataset as representing dying or damaged cells. The remaining clusters were annotated based on known marker genes.

### RNA velocity through dynamical modeling

We analyzed the RNA velocity to investigate developmental trajectories by recovering directed dynamic gene information through splicing kinetics. Information like clustering and UMAP coordinates were retrieved from the Scanpy analysis. The pre-processing and downstream analysis were performed with scVelo using default parameters ^20^. Splice variants and cells were filtered, normalized, and logarithmized with the function scv.pp.filter_and_normalize (parameters: min_cells=3, min_counts=200, min_shared_counts=20, n_top_genes=5500). The moments based on the connectivities were calculated with 30 PCAs and 30 neighbors in the next step. After recovering the dynamics, the latent time was calculated and the velocity was calculated as a dynamical model.

### Integration of datasets from different sequencing approaches

For integration and correction of datasets from different sequencing runs (**Fig. 3, Fig. S1, Fig. S2, and Fig. S7**), we applied bbknn to the datasets (neighbors_within_batch=20, n_pcs=30, trim=0, copy=True). We then reclustered the cells with the Leiden algorithm at a resolution of 0.8^75,76^.

### Transcription factor enrichment analysis

The identified cluster-specific DDGs or DEGs were entered in the ChEA3 web tool^77^ and the mean rank was plotted using R.

### Pathway and Gene Ontology enrichment

DEGs were filtered by their unique expression over all clusters (standard deviation above 0.5) and an expression value above 0.5. For the GO term enrichment, the R package enrichR was used with the “GO Biological Process 2018” database and plotted in R^78,79^.

### CellphoneDB

The count matrix and cluster annotation were exported from scanpy, imported into R, and processed as recommended by the authors^80^. Cell-cell interactions were selected by the highest mean score and lowest p-value.

### NicheNet

As Target gene input, top 300 DEGs have been used. The calculation was done in R, converting the anndata element into a Seurat object. The process was performed as recommended by the authors^60^.

### Software specifications

The scRNA-seq alignment was run in CellRanger version 3.0.1 and the analyses were run in python 3.7.4 with Scanpy API version 1.4.4 or 1.5.1, anndata version 0.6.22 or 0.7.4, umap version 0.3.10, numpy version 1.17.4, scipy version 1.5.1, pandas version 0.25.3 or 1.0.5, scikit-learn version 0.22, statsmodels version 0.10.1, python-igraph version 0.7.1, scvelo version 0.2.1, matplotlib version 3.2.1, seaborn version 0.9.0, loompy version 3.0.6, XlsxWriter version 1.2.6, bbknn version 1.3.6 and scrublet version 0.2.1.

The plots of TFEA and GO term analysis were generated in RStudio with R version 3.6.0 with the usage of the R packages enrichR_3.0, ggpubr_0.4.0, ggplot2_3.3.3, stringr_1.4.0, EBImage_4.32.0, and bioimagetools_1.1.5.

NicheNet analysis was performed using following package versions: xlsx_0.6.5, ggpubr_0.4.0.999, cowplot_1.1.1, RColorBrewer_1.1-2, circlize_0.4.13, forcats_0.5.1, stringr_1.4.0, dplyr_1.0.7, purrr_0.3.4, readr_2.1.1, tidyr_1.1.4, tibble_3.1.0, ggplot2_3.3.5, tidyverse_1.3.1, SeuratObject_4.0.4, Seurat_4.0.2, nichenetr_1.0.0, gridBase_0.4-7, and ComplexHeatmap_2.6.2

## Supporting information

lists of DEGs, DDGs, ligands, and antibodies

## Data availability

Raw data will be available via Gene Expression Omnibus.

## Code availability

The code for scRNA-seq analysis will be available on github.

## Acknowledgements

This work is supported by the Helmholtz Pioneer Campus, and ERC (Consolidator Grant Number 772646), We thank Thomas Walzthöni for bioinformatics support provided at the Bioinformatics Core Facility, Institute of Computational Biology, Helmholtz Zentrum München.

## Author contributions

S.R., M.A., and M.Meier. designed the study. S.R., M.A., C.B., M.Marder., and M.Meier. executed the biological experiments. S.R., M.A. did the imaging and image analysis. S.R. S.W. performed the scRNA-seq analysis. S.U., F.T. and M.Meier supervised the study. The manuscript was written by S.R., and M.Meier. All authors corrected and approved the paper.

