## supplementary tables and figures for "Single-cell characterization of neovascularization using hiPSC-derived endothelial cells in a 3D microenvironment"

**Table S1 | Cell sample and sequencing statistics.**

| sample ID | condition | dead cells / % | mean number of genes per cell | mean number of sequencing read counts | Number of analyzed cells |
|---:|---:|---:|---:|---:|---:|
| **day 0** | 3D suspension culture | 17.4 | 4440 | 22361 | 6411 |
| **day 3** | 3D suspension culture | 36.5 | 4132 | 19660 | 4549 |
| **day 6** | 3D suspension culture | 19.1 | 3644 | 12080 | 4807 |
| **day 9_1** | 3D suspension culture | 25.0 | 4051 | 15200 | 6425 |
| **day 9_2** | 3D suspension culture | 30.8 | 4388 | 19199 | 3011 |
| **day 12** | 3D suspension culture | 16.1 | 5354 | 24817 | 4114 |
| **day 12** | 3D Matrigel | 4.3 | 3835 | 12165 | 13530 |
| **day 18** | reaggregated 3D suspension culture | 17.0 | 4083 | 12869 | 4396 |
| **day 18** | 3D Matrigel | 12.3 | 4807 | 19542 | 6462 |
| **day 18** | 3D Matrigel + ascorbic acid | 7.6 | 4673 | 19369 | 8743 |

**Table S2 | Culturing conditions of the differentiation in 2D and 3D.**

| | **day of differentiation** | | **this study** | **McCracken et al.** | **day of differentiation** |
|---|---|---|---|---|---|
| **lateral mesoderm induction** | 0-3 | media | N2B27 | N2B27 | 1-4 |
| | | BMP4 | 25 ng/ml | 25 ng/mL | |
| | | CHIR99021 | 7.5 µM | 7 µM | |
| **endothelial induction** | 3 to 7 | media | StemPro-34 | StemPro-34 | 4 |
| | | VEGF-A | 200 ng/ml | 200 ng/ml | |
| | | Forskolin | 2 µM | 2 µM | |
| **late stage** | from 8 on | media | StemPro-34 | EGM-2 | until 8 |
| | | VEGF-A | 30 ng/ml | 50 ng/mL | |
| | | FGF-2 | 30 ng/ml | | |
| | | human AB serum | | 1% | |

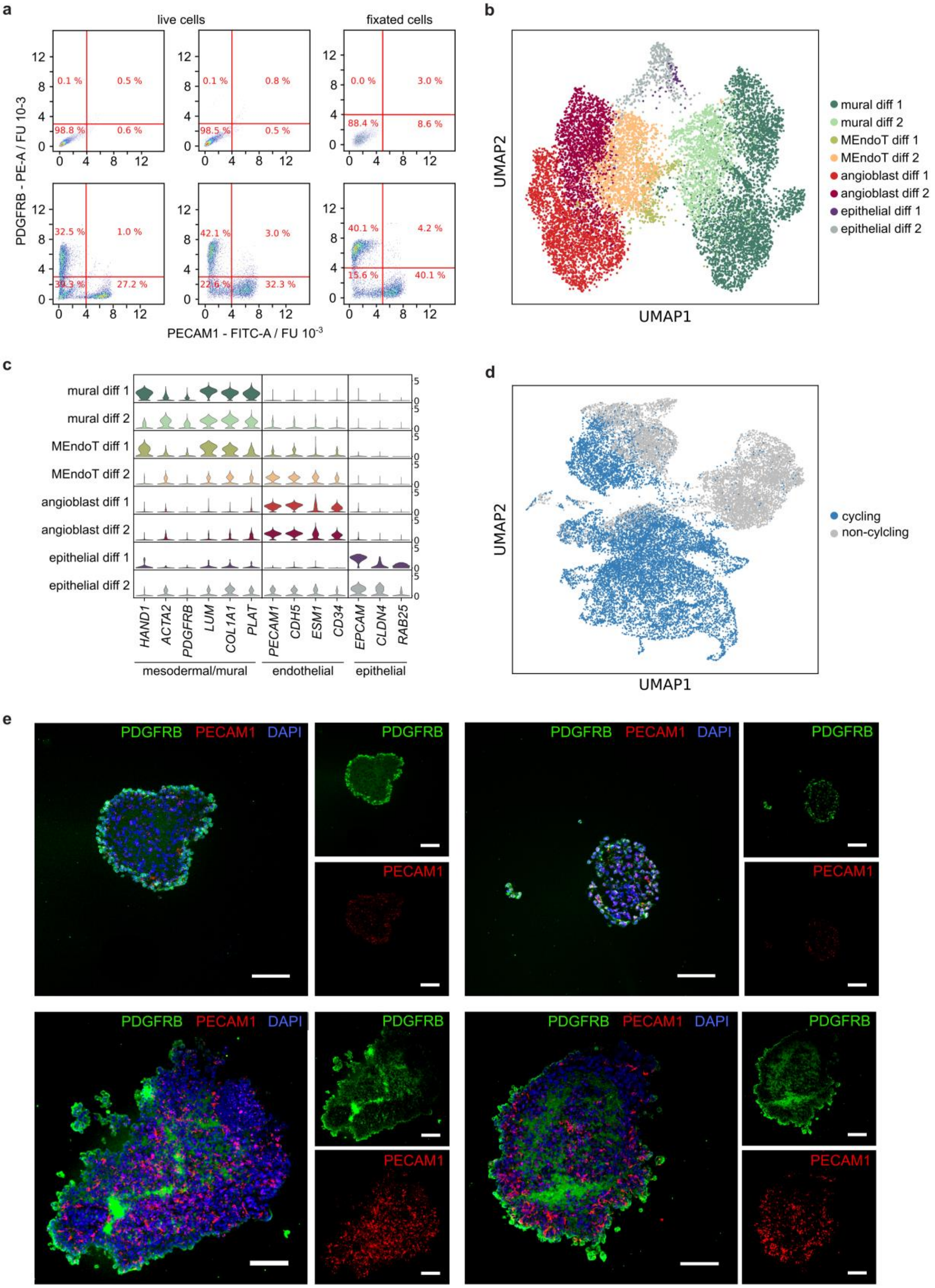
a
live cells
fixated cells
PDGFRB - PE-A / FU 10-3
PECAM1 - FITC-A / FU 10-3
0.1 %
0.5 %
98.8 %
0.6 %
0.1 %
0.8 %
98.5 %
0.5 %
0.0 %
3.0 %
88.4 %
8.6 %
32.5 %
1.0 %
39.3 %
27.2 %
42.1 %
3.0 %
22.6 %
32.3 %
40.1 %
4.2 %
15.6 %
40.1 %
b
UMAP1
UMAP2
mural diff 1
mural diff 2
MEndoT diff 1
MEndoT diff 2
angioblast diff 1
angioblast diff 2
epithelial diff 1
epithelial diff 2
c
HAND1
ACTA2
PDGFRB
LUM
COL1A1
PLAT
PECAM1
CDH5
ESM1
CD34
EPCAM
CLDN4
RAB25
mesodermal/mural
endothelial
epithelial
d
UMAP1
UMAP2
cycling
non-cylcling
e
PDGFRB PECAM1 DAPI
PDGFRB
PECAM1

**Fig. S1 | Reproducibility of the endothelial differentiation in a 3D suspension culture. a**, Flow cytometric analysis of three independent EC differentiation experiments. Cells were stained with PECAM1-FITC and PDGFRB-PE antibodies on day nine of differentiation. Upper left, unstained cell sample control; upper center and right, isotype control of the first EC differentiation experiment; lower row, three independent EC differentiation experiments. **b**, UMAP plot of transcriptomes from cells differentiated until day nine within two independent differentiation experiments and colored by annotated cell types (1, 2: mural, 3, 4: angioblast, 5, 6: MEndoT, 7, 8: epithelial, uneven numbers refer to first even to second differentiation). 6618 cells were analyzed in the first and 5035 in the second sequencing experiment. **c**, Violin plot of common cell type marker genes for annotation of the clusters. The density distribution indicates the normalized cluster mean expression. **d,** UMAP plot of the scRNA-seq dataset from experiment 1 (see also **Fig. 1** of the main text), where cells in G2, S, and M phase were colored in blue. **e**, Immunofluorescence images of sections of 3D suspension culture aggregates from day six (upper row) and day nine (lower row) stained for DAPI (blue), PDGFRB (green), and PECAM1 (red). Scale bar denotes 100 µm.

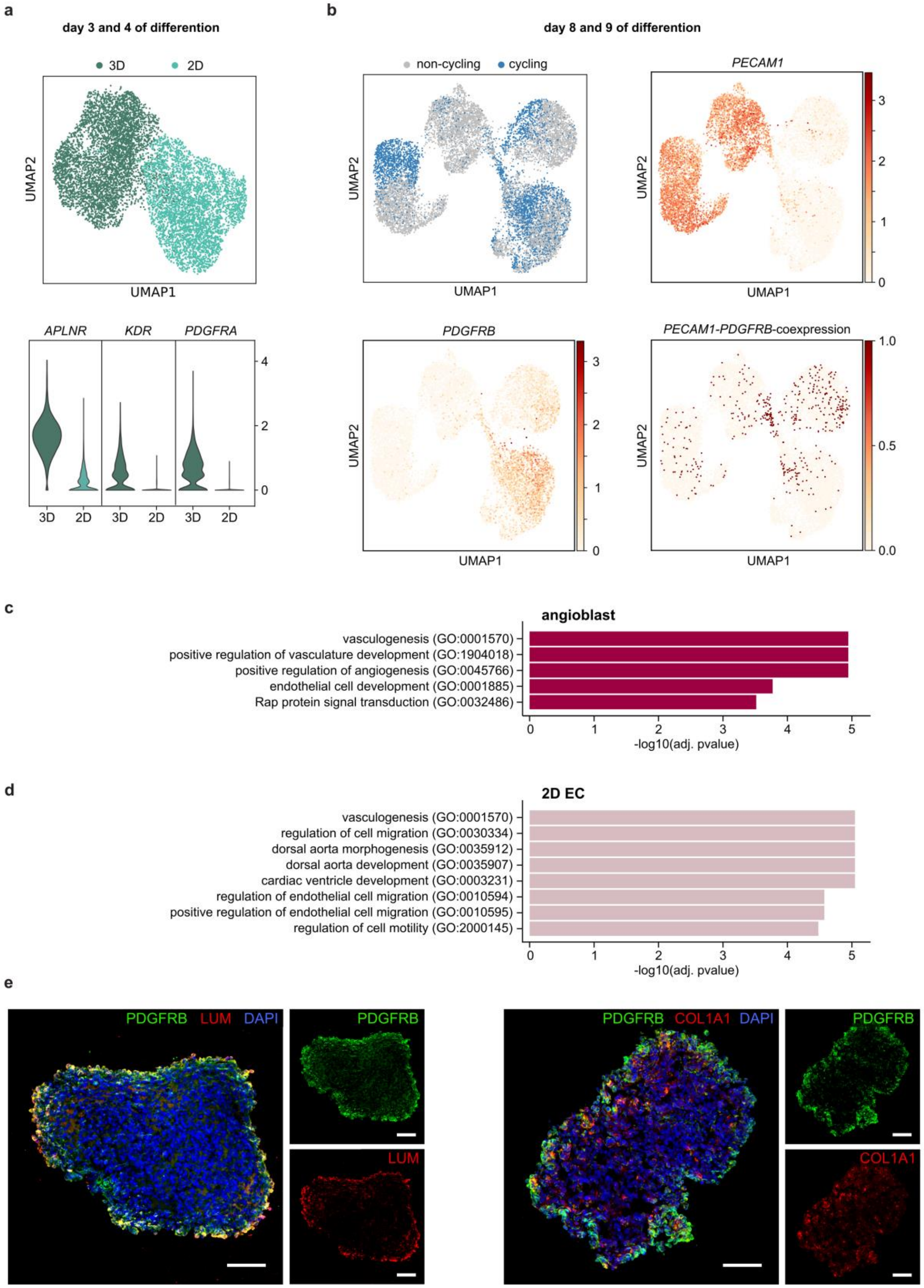
a
day 3 and 4 of differention
3D
2D
UMAP2
UMAP1
APLNR
KDR
PDGFRA
b
day 8 and 9 of differention
non-cycling
cycling
PECAM1
PDGFRB
PECAM1-PDGFRB-coexpression
c
angioblast
vasculogenesis (GO:0001570)
positive regulation of vasculature development (GO:1904018)
positive regulation of angiogenesis (GO:0045766)
endothelial cell development (GO:0001885)
Rap protein signal transduction (GO:0032486)
-log10(adj. pvalue)
d
2D EC
vasculogenesis (GO:0001570)
regulation of cell migration (GO:0030334)
dorsal aorta morphogenesis (GO:0035912)
dorsal aorta development (GO:0035907)
cardiac ventricle development (GO:0003231)
regulation of endothelial cell migration (GO:0010594)
positive regulation of endothelial cell migration (GO:0010595)
regulation of cell motility (GO:2000145)
-log10(adj. pvalue)
e
PDGFRB LUM DAPI
PDGFRB
LUM
PDGFRB COL1A1 DAPI
COL1A1

**Fig. S2 | Comparison of the EC differentiation in 2D and 3D cell culture formats. a**, Upper: UMAP plot of sc-transcriptomes acquired at days three and four (McCracken et al.) of the EC differentiation. Lower: Violin plots of mesenchymo-angioblast marker genes corresponding to the UMAP plot above. **b**, UMAP plots of the sc-transcriptomes acquired on day nine and day eight (McCracken et al.) of the EC differentiation. Upper left: Blue colored dots represent cells with gene expression relating to S, G2 or M-phase. UMAP plots colored by gene expression of *PECAM1* (upper right), *PDGFRB* (lower left) and their co-expression (lower right). **c** and **d**, Gene ontology term analysis based on DEGs from the sc-transcriptomes of ECs derived from stem cells cultured in a 2D monolayer and 3D suspension culture. **e**, Immunofluorescence images of sections of 3D suspension culture aggregates from day nine stained for DAPI, PDGFRB and either LUM (left) or COL1 (right). Scale bar denotes 100 µm.

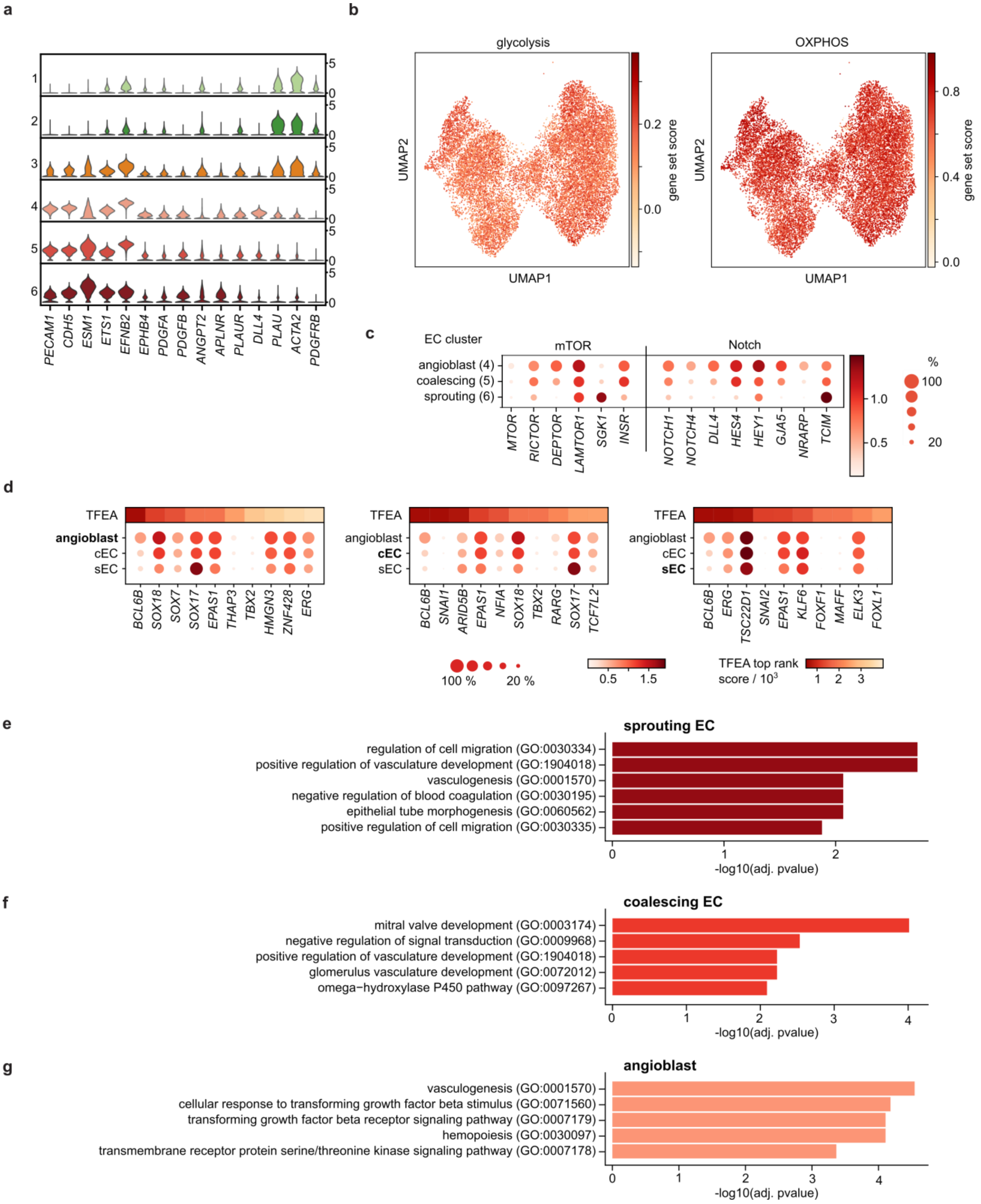

**Fig. S3 | Single-cell transcriptomes analysis of ECs and mural cells in Matrigel and 3D suspension culture. a,** Violin plot of representative marker genes used to assign the cell clusters in the UMAP plot in **Fig 4**. The density distribution indicates the normalized cluster mean expression. **b**, Enriched analysis of glycolysis (left), and OXPHOS (right) genes based on mean expressions. The plot showed that enzymes within the metabolic pathways were not differentially regulated in the angioblast, coalescing, or sprouting EC transcriptional state. **c**, Dot plot of genes within the mTOR and Notch pathway regulated during the cell state transition from angioblast to coalescing ECs and sprouting ECs. **d**, Transcription factor enrichment analysis based DEGs the angioblast (left), coalescing (middle), and sprouting (right) EC cluster. DEGs between the EC clusters were filtered for expression and standard deviation before TFEA. TFEA scores are represented in a color code, whereas the mean expression levels of the corresponding TFs as dot plots. **e**-**g**, Gene ontology term analysis based on DEGs from the sc-transcriptomes of angioblast to coalescing EC and sprouting ECs.

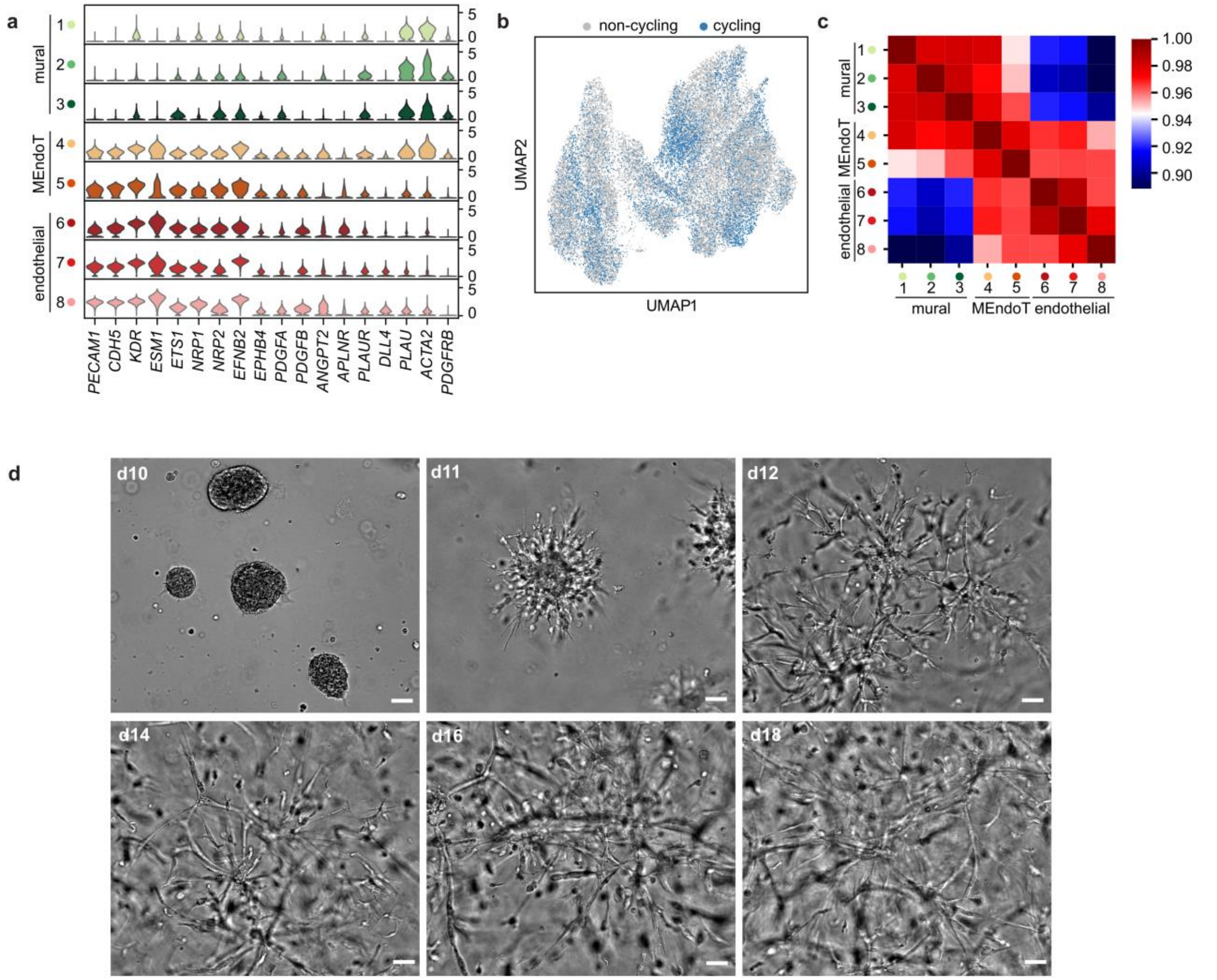

**Fig. S4 | Stem-cell derived endothelial cells form microvessels in Matrigel culture with and without ascorbic acid. a**, Violin plot of marker genes used to assign the cell cluster. The density distribution indicates the normalized cluster mean expression. **b**, UMAP plot of sc-transcriptomes acquired from cells in the Matrigel cultures on days 12 and 18. Blue colored dots represent cells expressing genes relating to S, G2 or M-phase. **c**, Cell cluster Pearson correlation plot. **d**, Bright-field images of the Matrigel culture along the timeline of day 10 to 18. Scale: 50 µm.

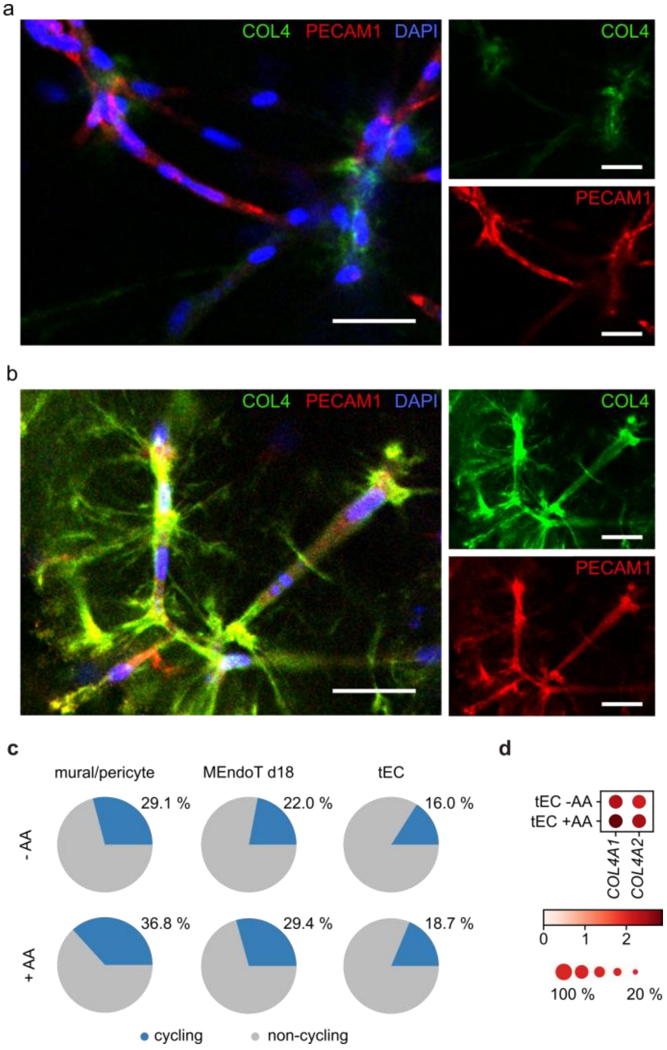

**Fig. S5 | Immunofluorescence images of microvessels at day 18 in the Matrigel culture with and without the addition of ascorbic acid. a**, Immunofluorescence image of Matrigel culture on day 18 without and **b**, with AA addition to the media, where COL4, DAPI and PECAM1 were counterstained. **c**, Cluster-specific proliferation rate for day 18 specimen with and without ascorbic acid. **d**, Expression levels of the collagen IV chains between the condition with and without AA inside the tEC cluster. Color intensity and dot size denote the normalized cluster mean expression level and fraction of cell expression for the corresponding gene, respectively.

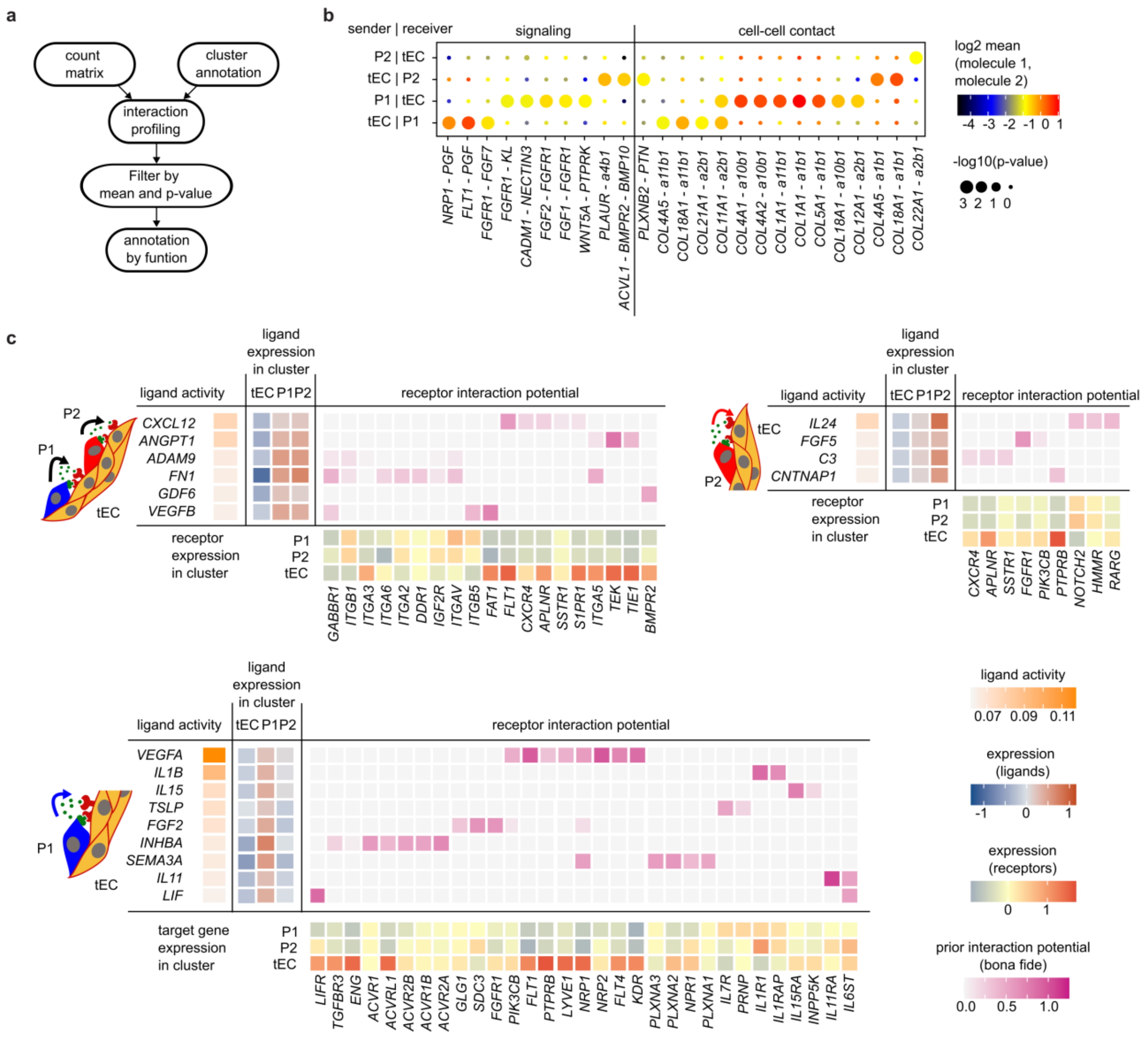

**Fig. S6 | Ligand-receptor interaction inferred by CellphoneDB and NicheNet. a**, Schematic of CellphoneDB procedure. **b**, Dot plot with the highest calculated ligand-receptor links between tubulogenic ECs (tEC) and the two pericyte clusters (P1/2), respectively. The circle size indicates the p-values and the color of the mean expression levels. **c**, NicheNet predicted receptor interaction potential in the receiver cells for the ligands send by either tubulogenic ECs (tEC) or the two pericytes.

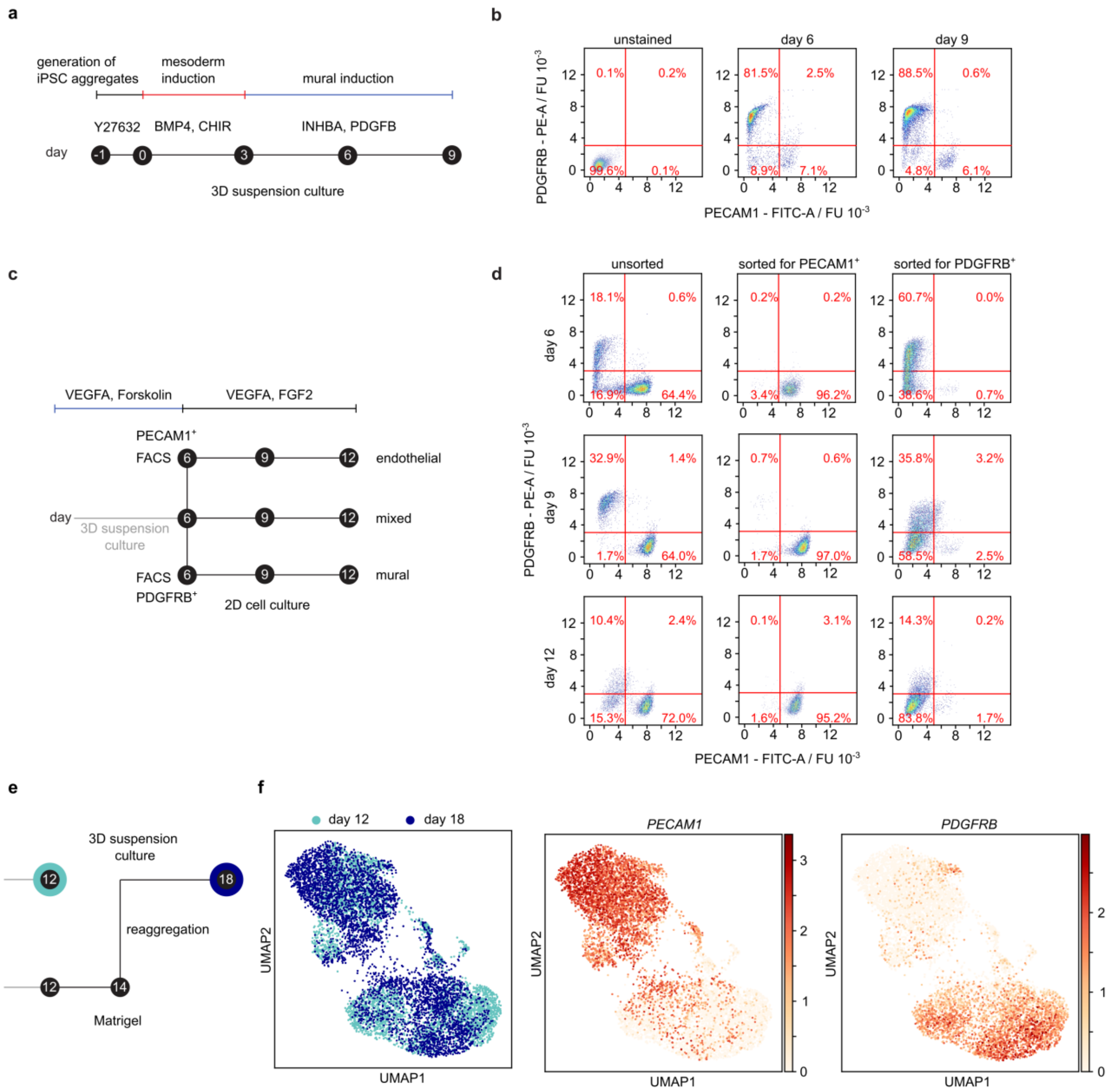

**Fig. S7 | Cell plasticity of ECs shown upon various culturing conditions. a**, Schematic of mural cell differentiation in 3D suspension culture, **b**, Flow cytometry analysis of days 6 and 9 of the differentiation. **c**, Experimental overview of cell sorting on day 6 of the EC differentiation for endothelial and mural marker PECAM1 and PDGFRB and the three different culturing conditions. **d**, Flow cytometry analysis of sorted and unsorted cell populations from days 6, 9, and 12 cultured under EC differentiation media. **e**, Schematic of the experimental timeline and applied culture conditions. First, microvessels were formed upon transfer of 3D suspension culture into Matrigel on day 10 of differentiation. Second, the microvessels were disaggregated by harvesting single-cell solution on day 14. Third, single cells were reaggregated and cultured as 3D suspension culture until day 18. **f**, UMAP plots of sc-transcriptomes measured from 3D suspension cultures cultured to day 12 of differentiation (cyan) and reaggregated 3D suspension-cultured to day 18 (dark blue, left), colored for the gene expression of the EC marker *PECAM1* (middle) and mural cell marker *PDGFRB* (right).
